# A Biophysical Basis for Learning and Transmitting Sensory Predictions

**DOI:** 10.1101/2022.10.31.514538

**Authors:** Salomon Z. Muller, LF Abbott, Nathaniel B. Sawtell

## Abstract

Homeostatic (anti-Hebbian) forms of synaptic are effective at eliminating “prediction errors” that signal the differences between predicted and actual sensory input. However, such mechanisms appear to preclude the possibility of transmitting the resulting predictions to downstream circuits, severely limiting their utility. Using modeling and recordings from the electrosensory lobe of mormyrid fish, we reveal interactions between axonal and dendritic spikes that support both the learning *and* transmission of predictions. We find that sensory input modulates the rate of dendritic spikes by adjusting the amplitude of backpropagating axonal action potentials. Homeostatic plasticity counteracts these effects through changes in the underlying membrane potential, allowing the dendritic spike rate to be restored to equilibrium while simultaneously transmitting predictions through modulation of the axonal spike rate. These results reveal how two types of spikes dramatically enhance the computational power of single neurons in support of an ethologically relevant multi-layer computation.

## Introduction

The synaptic plasticity associated with learning is typically Hebbian or similar to Hebbian in that changes induced by plasticity lead to further plasticity (Abbott and Nelson, 2000; Caporale and Dan, 2008). This instability is what drives learning-related changes in neural responses, but it must be countered by some form of synaptic regulation that keeps synaptic strengths bounded (Miller and Mackay, 1994). In contrast, homeostatic forms of plasticity compensate for changing inputs by returning neural response back to an equilibrium point and hence do not require additional regulation (Bell et al., 1997c; Roberts and Bell, 2002; Turrigiano, 2017). Although the intrinsic stability of homeostatic plasticity is a powerful feature, it has generally been assumed to limit the utility of this form of plasticity for the transmission of learned signals.

Homeostatic forms of synaptic plasticity have been studied extensively in cerebellum-like sensory structures, where they play a critical role in the adaptive cancellation of self-generated sensory stimuli (Bell et al., 1997a; Bell et al., 2008). Cerebellum-like sensory structures combine input from peripheral sensory receptors--e.g. electroreceptors in the case of the electrosensory lobe (ELL) of teleost fish and auditory nerve fibers in the case of the mammalian dorsal cochlear nucleus (DCN)--with massive input from a mossy fiber-granule cell-parallel fiber system similar to that of the cerebellum. Granule cells convey a rich variety of information including: motor corollary discharge, proprioception, input from other sensory modalities, and feedback from higher processing stages within the same sensory modality. Numerous lines of evidence from *in vitro* and *in vivo* recording studies indicate that homeostatic (anti-Hebbian) forms of plasticity at parallel fiber synapses drive postsynaptic firing to a constant equilibrium rate by forming negative images of sensory input patterns that are predictable based on granule cell input (Bell, 1981; Bell et al., 1997c; Bodznick et al., 1999; Harvey-Girard et al., 2010; Kennedy et al., 2014). While computational models based on these results elegantly explain how homeostatic plasticity cancels predictable, self-generated sensory input within individual neurons (Kennedy et al., 2014; Roberts and Bell, 2000), there is a disconnect between these models and the actual circuitry and function of cerebellum-like structures. For both the ELL of mormyrid fish and the mammalian DCN, the major site of anti-Hebbian plasticity is at an intermediate stage of processing at parallel fiber synapses onto interneurons that inhibit output neurons (Bell et al., 1997c; Fujino and Oertel, 2003; Meek et al., 1996; Tzounopoulos et al., 2004). Homeostatic plasticity that maintains postsynaptic firing rate at a constant equilibrium rate would seemingly preclude interneurons from transmitting learned signals to the critical output stage of the network. This problem is not specific to the ELL, but would confront any system that relies on homeostatic plasticity to predict sensory input (Hertag and Sprekeler, 2020; Keller and Mrsic-Flogel, 2018).

A recent study of interneurons in the mormyrid ELL, known as medium ganglion (MG) cells, shows that this seeming paradox is resolved by the separate axonal and dendritic spikes found in these neurons (Muller et al., 2019). Consistent with prior experiments and modeling, the effects of self-generated sensory input on dendritic spike rate were cancelled by negative images. Unexpectedly, however, negative images were encoded in the axonal spike rate of MG cells (Figure 1A). The axonal transmission of negative images supports multi-layer computation in the ELL by enabling homeostatic plasticity at intermediate layer synapses between granule cells and MG cells to aid in cancelling predictable sensory input at the output layer of the ELL. However, these findings are puzzling from a biophysical standpoint: because granule cell input is sculpted by homeostatic plasticity to cancel predictable sensory input, the net input to the MG cell should be constant, and both dendritic and axonal spike rates should be unmodulated.

**Figure 1.**
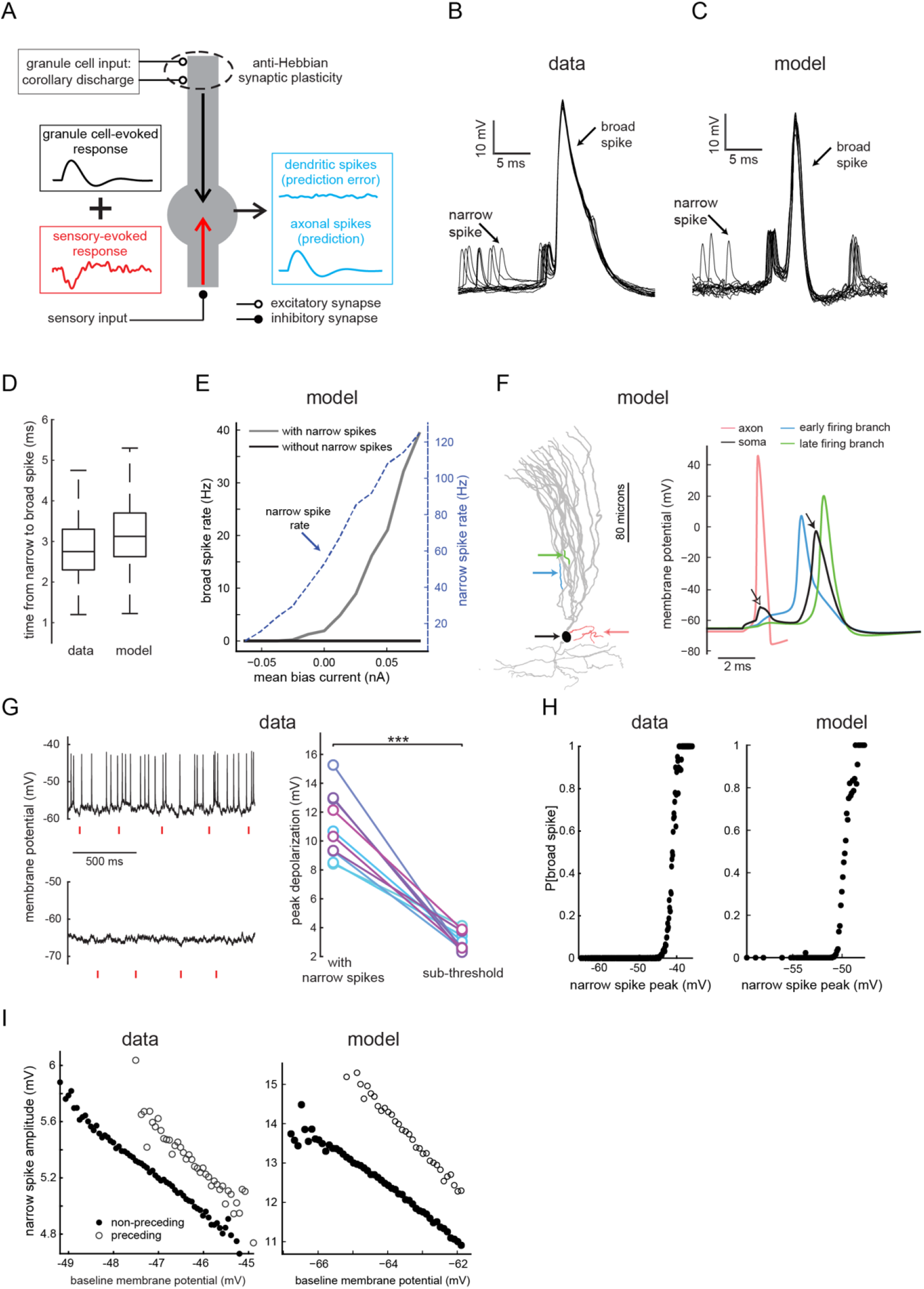
Backpropagating narrow spikes evoke broad spikes. (**A**) Schematic of negative image formation and transmission in MG cells. Electrosensory input containing both self-generated signals (arising from fish’s own electric organ discharge) and external signals (arising from prey) (red) is relayed to the basilar dendrites of MG cells via inhibitory and excitatory interneurons (not shown). Anti-Hebbian plasticity at granule cell synapses creates negative images of self-generated signals (black trace) based on motor corollary discharge signals (as well as other types of information) conveyed by granule cells. Granule cell input cancels the effects of self-generated sensory input on dendritic spikes (blue trace, top) but modulates axonal spikes (blue trace, bottom). (**B-C**) Overlaid intracellular voltage traces from an example MG cell recorded *in vivo* (**B**), (and see Figure 1-figure supplement 1D) and the model cell (**C**). (**D**) Interval between peaks of narrow and broad spikes in recorded (n=17) and model MG cells. (**E**) Effect of eliminating narrow spikes on broad spike firing in the model. Narrow-spike F-I curve is also shown. (**F**) Left, neurolucida reconstruction of an MG cell used to build the multi-compartment model. Arrows indicate the sites of the membrane voltage recordings depicting the process of broad spike initiation (right). Open and filled arrows indicate somatically recorded narrow and broad spikes, respectively. Voltage trace from the axon is truncated for clarity (omitted portion shows that broad spikes trigger an additional axonal spike). (**G**) Left, membrane potential fluctuations in an MG cell recorded with no bias current (top) and with hyperpolarizing bias current to prevent narrow spiking (bottom). Red lines indicate the times of the fish’s electric organ discharge command. Right, peak depolarization amplitudes (relative to baseline) are substantially larger with narrow spikes intact (n=10 p< 0.001). (**H**) Left, example MG cell recording illustrating the relationship between broad spike probability and the peak of the narrow spike immediately preceding the broad spike. Additional examples are shown in Figure 1-figure supplement 1F-G. Right, same display for the model cell. (**I**) Narrow spike amplitude depends on the baseline membrane potential (i.e. the point from which the spike arises) but for any given baseline membrane potential, narrow spikes that precede broad spikes have, on average, a larger amplitude. Left panel shows one example MG cell (and see Figure 1-figure supplement 1H-K) and right panel shows results from the model cell. Each circle represents the average amplitude for the given baseline membrane potential.

Here we combine *in vivo* intracellular recordings with biophysical modeling to show that this puzzle is resolved at the biophysical level by interactions between inhibitory synaptic input, backpropagating axonal action potentials, and dendritic spikes. Specifically, we demonstrate: (1) that backpropagating axonal action potential evoke dendritic spikes in MG cells and (2) that sensory inhibition (or disinhibition) can selectively modulate the rate of dendritic spikes by controlling backpropagating spike amplitude. These interactions allow homeostatic plasticity to maintain a constant dendritic spike rate by enforcing cancellation while simultaneously inducing modulations in axonal spike rate that transmit sensory predictions. Our modeling work is based on a multi-compartment neuronal model, but the basic results are recapitulated in a model with only somatic and axonal compartments. Thus the mechanism we describe does not require dendritic computation, but relies instead on an electrotonically distant spike initiation site in the axon—a common feature of neurons that, to our knowledge, has not been previously connected to learning.

## Results

### Dendritic spikes are triggered by backpropagating axonal spikes

MG cells fire two types of sodium channel-dependent action potentials known as broad and narrow spikes (Figure 1B). Broad spikes are likely initiated in the proximal apical dendrites, have a high threshold, and are emitted at low rates, while narrow spikes are likely initiated in the axon, have a low threshold, and are emitted at high rates (Bell et al., 1997b; Engelmann et al., 2008; Grant et al., 1998). Broad spikes induce long-term depression at granule-MG cell synapses (Bell et al., 1997c; Han et al., 2000). Given their critical role in plasticity induction, we sought to determine what factors control broad spike firing *in vivo*. Confirming prior studies (Bell et al., 1997b; Grant et al., 1998; Sawtell et al., 2007), we found that broad spikes are invariably preceded by a narrow spike at a characteristic interval of ~3 ms (Figure 1B,D and Figure 1-figure supplement 1A-B). This observation, by itself, does not indicate that narrow spikes play a causal role in evoking broad spikes, as preceding narrow spikes could arise simply because broad spikes have a higher threshold than narrow spikes. To further evaluate this question, we examined a previously developed multi-compartment MG cell model_that recapitulates critical features of MG cell responses described above (Figure 1A, *right*) (Muller et al., 2019). This model is reduced, containing a minimal set of voltage-gated and synaptic conductances (Materials and methods), which makes it amenable to detailed analysis. The model is tuned to produce observed broad and narrow spike rates, but no fine-tuning of parameters is required to produce the results we report. In fact, as we show later, the basic effects can be reproduced in a further reduced model.

When input currents were adjusted in the model to evoke the ~50 Hz narrow spike firing and ~2 Hz broad spike firing seen *in vivo*, broad spikes in the model cell were always preceded by a narrow spike at an interval of ~3 ms (Figure 1C-D and Figure 1-figure supplement 1B). Blocking narrow spikes by turning off active conductances in the axonal compartment abolished broad spike firing over a range of input strengths (Figure 1E), while injecting a brief spike-like depolarizing current into the soma (with active conductances in the axon turned off) evoked broad spikes after a similar delay (Figure 1-figure supplement 1C). These results confirm a causal role for narrow spikes in evoking broad spikes in the model. Monitoring the voltage at various locations revealed that even though axonal depolarization resulting from the narrow spike is highly attenuated by the time it reaches the soma (Figure 1F, *open arrowhead*), it nevertheless spreads passively into the proximal apical dendrites where it activates voltage-gated sodium and potassium channels to evoke a local dendritic spike (Figure 1F, *blue*). Depolarization from the local dendritic spike then propagates into other apical branches leading to additional spike initiations at multiple sites throughout the apical dendrite. These local dendritic spikes sum to produce a broad somatic spike after a delay of several milliseconds from the triggering narrow spike (Figure 1F, *filled arrowhead* and Video). Characteristics of putative apical dendritic MG cell recordings *in vivo* are consistent with the model; narrow spikes are smaller and broad spikes are narrower in comparison with somatic recordings (Figure 1-figure supplement 1D-E).

Several lines of evidence suggest a causal role for backpropagating narrow spike in evoking broad spikes *in vivo* through a process similar to that described in the model. First, eliminating narrow spikes with hyperpolarizing current or a sodium channel blocker in the recording pipette revealed that peak somatic depolarization due to narrow spikes is much greater than that due to subthreshold input alone (Figure 1G). Second, *in vivo* (as in the model), the probability of evoking a broad spike depends strongly on the membrane potential at the peak of the recorded narrow spike (Figure 1H and Figure 1-figure supplement 1G). Third, *in vivo* (as in the model), narrow spikes that immediately precede a broad spike not only arise from more depolarized potentials (as would be expected based on the higher threshold for broad spikes) but also exhibit larger amplitudes than narrow spikes not preceding a broad spike (Figure 1I and Figure 1-figure supplement 1H-K). It is difficult to see why this would be the case if the narrow spike were not causal. We also observed that the amplitude of backpropagating narrow spikes depends on the baseline membrane potential, an effect seen in other systems (Grace and Bunney, 1983), presumably due to the voltage-dependence of the membrane conductance.

### A biophysical model of negative image formation and transmission

We next examined how sensory input affects broad and narrow spike firing in the multicompartment model. *In vivo* studies have revealed two sub-classes of MG cells: BS-cells, in which the broad spike rate is decreased by sensory input, and BS+ cells, in which the broad spike rate is increased (presumably through dis-inhibition). While for simplicity we focus on modeling BS-cells, the mechanisms we describe can also explain responses in BS+ cells (Figure 2-figure supplement 1). For clarity, we consider constant sensory input, but we have verified that all the results we report apply to time-dependent sensory inputs matching those *in vivo* (Figure 2-figure supplement 3). Adding constant inhibitory input to basilar dendritic compartments potently reduces the broad spike firing rate with little effect on the rate of narrow spikes (Figure 2A, *inhibition*), consistent with prior *in vivo* recordings (Muller et al., 2019). Measuring membrane potential values in the somatic compartment of the model revealed that inhibition results in narrow spikes reaching less depolarized levels at their peaks (Figure 2B, *red* and Figure 2-figure supplement 2b), so that the broad spike threshold is rarely crossed (Figure 2B, *dashed line*). The increased conductance due to the inhibitory input reduces the peak membrane potential by attenuating the passive spread of the narrow spike from the axon initial segment, as seen in the small reduction in the amplitude of the narrow spike at the soma (~0.75 mv; Figure 2C, E, *red* and Figure 2-figure supplement 2C; Appendix 1). The effects of inhibitory input on the baseline membrane potential and the narrow spike rate are negligible because the narrow spike threshold (~-64 mV) is near the reversal potential for inhibition (−65 mV in the model) (Figure 3A).

**Figure 2.**
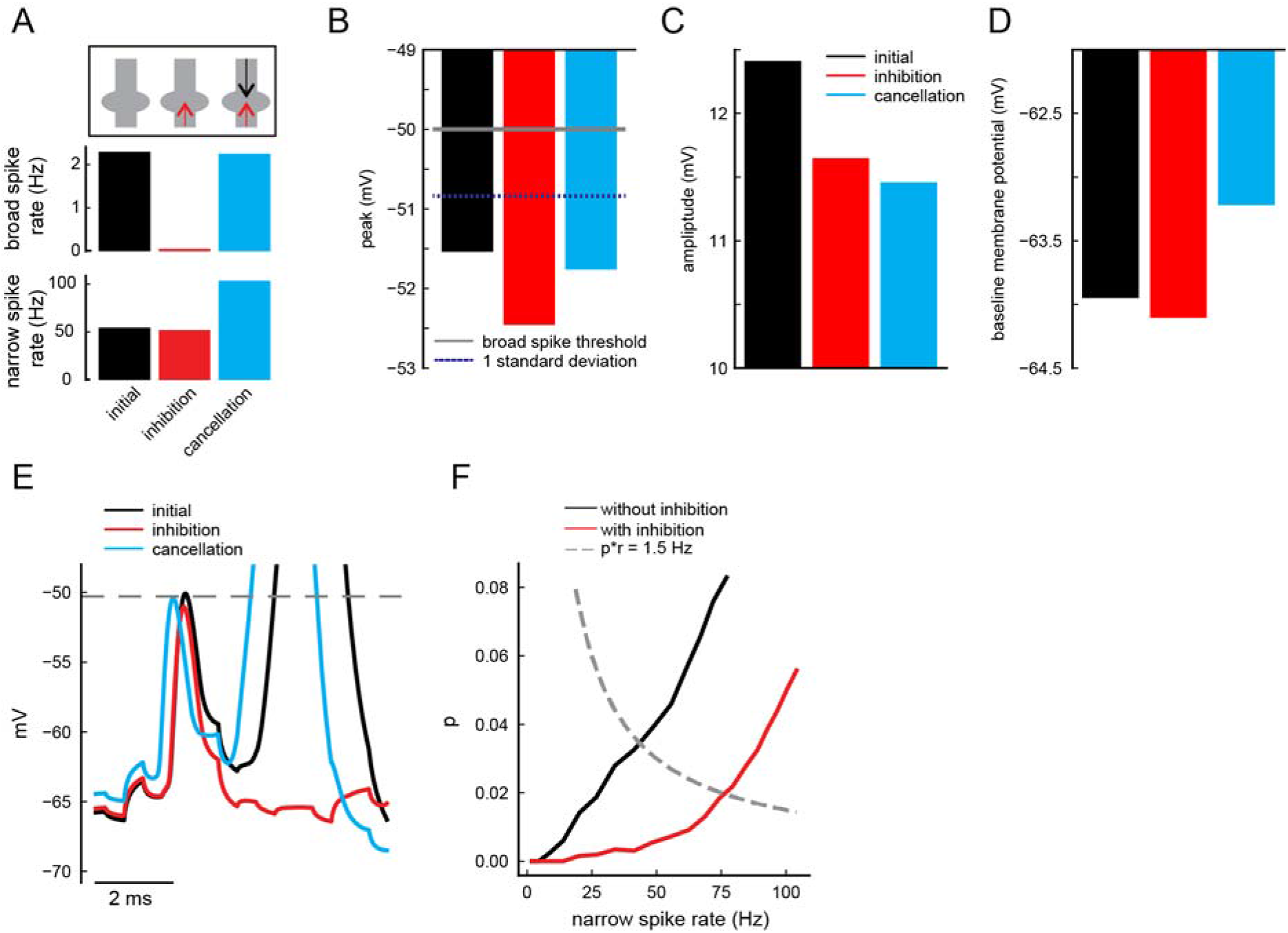
Biophysical model of negative image formation and transmission. (**A**) Narrow and broad spike rates under three conditions used to simulate the formation and transmission of negative images in the model (see main text). To simplify model analysis, we use step-like changes in sensory and corollary discharge input rather than simulating the temporal response profiles observed *in vivo* (Figure 2-figure supplement 3). This is equivalent to plotting the peak of the responses schematized in Figure 1A. (**B**) Peak membrane potential of backpropagating narrow spikes for the input conditions shown in (A). Gray line indicates the broad spike threshold and the distance from the gray line to the dashed blue line is one standard deviation from the mean (which is similar across all three conditions). (**C**) Backpropagating narrow spike amplitudes for the input conditions shown in (A). (**D**) Baseline membrane potentials for the input conditions shown in (A). (**E**) Example voltage traces from the model illustrating how membrane potential depolarization (cyan) allows narrow spikes to cross the threshold for evoking a broad spike (dashed line), despite the reduction in narrow spike amplitude due to inhibition (red). (**F**) Inhibition (red) reduces probability of evoking a broad spike (p), such that an increase in narrow spike rate is required to restore the broad spike rate to equilibrium (dashed line). This increase is proportional to the negative image. Equilibria for the two conditions are where the dashed and solid curves cross.

**Figure 3.**
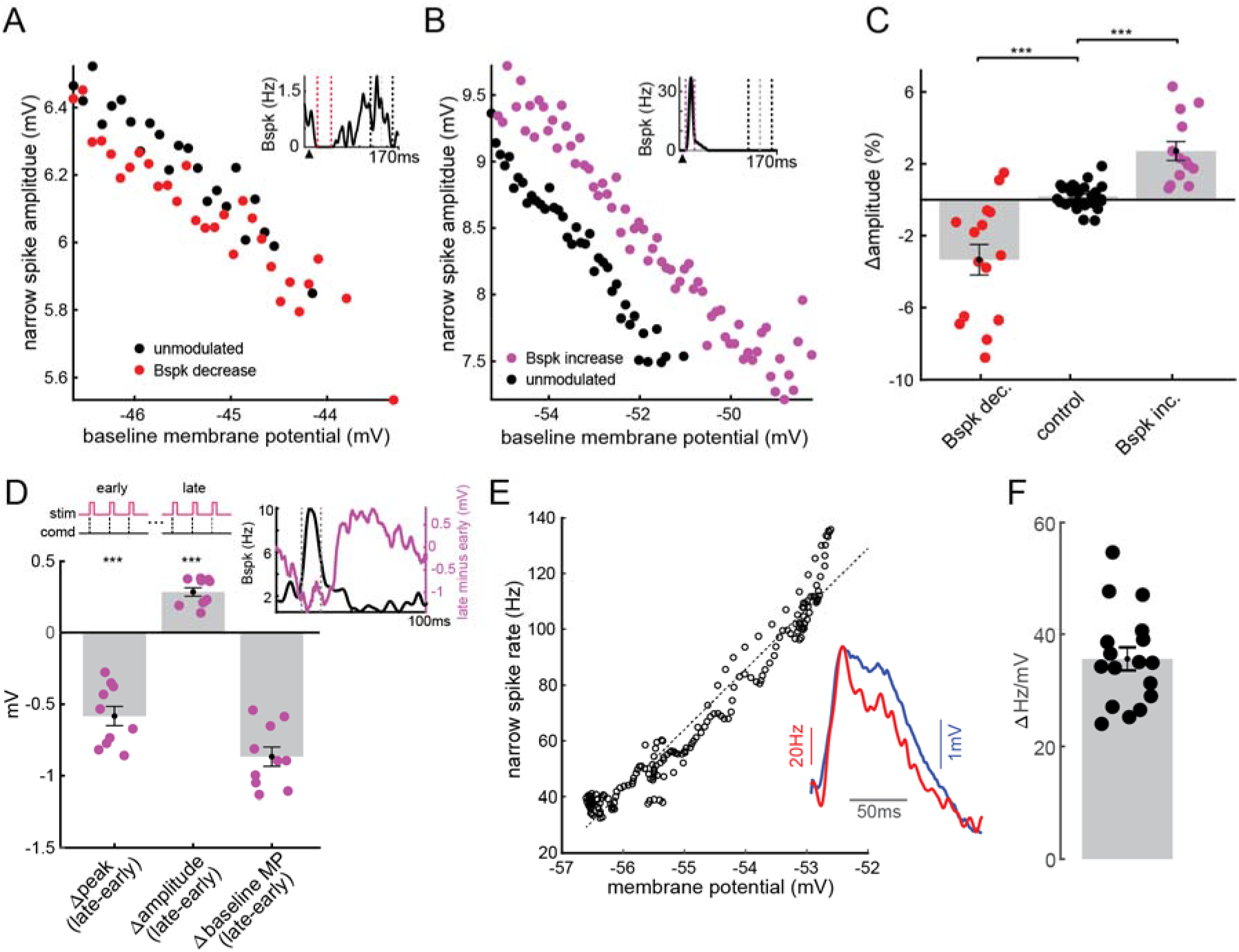
Negative image formation and transmission *in vivo*. (**A**) Example BS-MG cell illustrating a decrease in the backpropagating narrow spike amplitude in a time window when broad spike firing is transiently decreased by an electrosensory stimulus (red) compared to a window in which the broad spike rate is not modulated (black). Inset here and in (B) identifies these analysis windows. (**B**) Example BS+ MG cell illustrating an increase in the backpropagating narrow spike amplitude in a time window when broad spike firing was transiently increased by an electrosensory stimulus (magenta). (**C**) Summary of the effects of sensory stimuli on narrow spike amplitude across MG cells (n = 15 decrease, n = 13 increase, p<0.001). Middle bar (control) shows results of analysis comparing amplitudes in two windows in which broad spike rates were not modulated (the two windows are separated by the gray dashed line in insets A-B). (**D**) Changes in membrane potential at the peak of the narrow spike (Δpeak), narrow spike amplitude (Δamplitude), and the baseline membrane potential preceding narrow spikes (Δbaseline MP) during pairing (~4 minutes) of an electrosensory stimulus with the electric organ discharge motor command (comd) to induce negative image formation and sensory cancellation in BS+ cells (n = 10). Inset right, traces from an example cell illustrating the initial sensory-evoked increase in broad spike firing (black) along with the resulting change in the membrane potential, which forms an approximate negative image of the effects of the paired sensory input on broad spike firing (magenta). Inset left, illustration of the pairing paradigm. (**E**) Narrow spike rate versus membrane potential plotted for one example cell. Dashed line is the linear fit. Inset, trial-averaged membrane potential (with spike removed) and corresponding narrow spike rate for the same cell. (**F**) Average sensitivity of narrow spikes to membrane potential changes across MG cells (n=17) calculated based on the range of the curves shown in E.

Next, we examined the central question of how negative image formation affects broad and narrow spikes. The dynamics of anti-Hebbian spike timing-dependent plasticity acting on realistic granule cell inputs have been extensively characterized and modeled (Bell et al., 1997c; Kennedy et al., 2014; Roberts and Bell, 2000). Because the focus here is on the consequences of these well-characterized plasticity dynamics on narrow and broad spike firing (rather than on the plasticity mechanism itself), we simply reproduce the known effect of this plasticity in our model. In other words, we set the strengths of excitatory conductances onto apical dendrites to cancel the effects of inhibition on the broad spike rate_(Figure 2A, *cancellation*), which is precisely what the anti-Hebbian plasticity does. Sensory input temporarily lowers the broad spike rate but mimicking synaptic plasticity returns this rate back to its equilibrium value by restoring the membrane potential at the peak of the backpropagating narrow spike close to its baseline value (Figure 2B, *cyan* and Figure 2-figure supplement 2b). Critically, however, the reduction of the backpropagating narrow spike amplitude caused by inhibition is not reversed (Figure 2C, *cyan* and Figure 2-figure supplement 2C). Instead, restoration of the broad spike rate requires an additional depolarization of the underlying membrane potential that assures that the peak of the attenuated narrow spike reaches the threshold for broad spike firing (Figure 2D, E, *cyan* and Figure 2-figure supplement 2D). This baseline depolarization drives narrow spike firing, thereby transmitting the negative image to downstream neurons (Figure 2A, *cancellation*).

An equivalent computational explanation for this phenomenon can be constructed by expressing the broad spike rate, R_bs_, as the product of two factors, the probability of a narrow spike evoking a broad spike, p, and the rate of narrow spikes, r_ns_; R_bs_ = p·r_ns_. The factor p reflects the functional coupling between backpropagating narrow spikes and broad spikes (similar to the “safety factor” described in classical studies of initial segment-somatodendritic spike coupling (Coombs et al., 1957a, b; Fuortes et al., 1957; Renshaw, 1942). Sensory input selectively affects the broad spike rate by reducing the value of p. (While narrow spike peak voltage is the dominant factor affecting p (Figure 1H), other factors may also contribute (Figure 2-figure supplement 4)). Specifically, suppose that the broad spike rate R_bs_ = por_ns_ is at its equilibrium value in the absence of sensory input, with p = p_0_. Introducing inhibition due to sensory input reduces p causing the broad spike rate to decrease. Synaptic plasticity restores the broad spike rate by returning por_ns_ and thus R_bs_ back to its equilibrium value (Figure 2F, dashed line). However, through this process p is not restored to its previous value p_0_, but instead remains smaller than p_0_. Thus, the restoration of the broad spike rate requires a compensatory increase in r_ns_, the narrow spike rate (Figure 2F).

### Negative image formation and transmission *in vivo*

The model makes two key predictions regarding negative image generation and transmission that we tested *in vivo:* (1) sensory input modifies narrow spike amplitude and (2) synaptic plasticity restores the broad-spike rate in the presence of sensory input by modifying the baseline membrane potential rather than by reversing the effects of sensory input on the amplitude of backpropagating narrow spikes. Comparing narrow spike amplitudes in time windows when broad spike firing was modulated by an electrosensory stimulus versus control windows revealed that sensory stimuli that suppressed broad spiking reduced the amplitude of backpropagating narrow spikes, while stimuli that enhanced broad spiking increased this amplitude (Figure 3A-C and Figure 3-figure supplement 1). The former corresponds to BS-cells, while the latter corresponds to BS+ cells in which both the sign of the broad spike response to the sensory input and the negative image are reversed.

To test prediction 2 concerning the changes in baseline membrane potential, we examined narrow spike amplitudes during the learning of negative images induced by pairing an electrosensory stimulus with the motor command that discharges the electric organ (Bell, 1981). This analysis was only possible for BS+ cells because of the much faster time-course of cancellation in these cells (Muller et al., 2019). As expected, cancellation of sensory-evoked increases in broad spike firing was driven by a temporally-specific hyperpolarization of the underlying membrane potential (Figure 3D, inset). Importantly, sensory-evoked changes in narrow spike amplitude were not reversed as negative images formed, a critical feature for our model of negative image transmission (Figure 3D). In fact, the amplitude of backpropagating narrow spikes actually increased due to the prominent inverse correlation of the narrow spike amplitude and the baseline membrane potential (Figure 1I). This effect amplifies the mechanism identified in the model, leading to even more robust negative image transmission by narrow spikes (Figure 3-figure supplement 2). Defining ΔAmp as the change in narrow spike amplitude due to sensory input and S as the slope of the relationship between narrow spike amplitude and the baseline membrane potential, the negative image is equal to −ΔAmp/(1+S). Our data suggest a value for S of ~-4.1% (Figure 3-figure supplement 2) which corresponds to −0.62 mV for a typical 15 mV narrow spike recorded in the soma. Hence, the negative image generated by a 3% narrow spike amplitude change is expected to be 0.45mV/(1-0.62) = 1.2 mV. Based on measured dependence of narrow spike firing rate on membrane potential (Figure 3E, F), this amounts to a ~40 Hz change in narrow spike rate, consistent with the magnitude of negative images recorded *in vivo*.

### Axonal, but not dendritic, compartmentalization is required for MG cell function

The differential effect of sensory input on broad and narrow spikes might suggest that spatial targeting of synaptic inputs onto MG cells is essential for learning and transmitting negative images. We tested this by varying the location of the sensory input in the model. Surprisingly, inhibition onto the proximal apical dendrites (Figure 4A) or soma (Figure 4B) yielded similar model performance as inhibition onto basilar dendrites (Figure 2A). In both cases, sensory input robustly decreased broad spike firing with little effect on narrow spike firing (Figure 4A, B, *inhibition*), and the addition of excitatory input to the apical dendrites cancelled the effects of sensory input on the broad spike rate while simultaneously modulating narrow spike output (Figure 4A, B, *cancellation*). Furthermore, if a mixture of excitatory and inhibitory input is delivered to the basilar dendrites, narrow spike firing rate is also increased while broad spike firing decreased (Figure 4C), matching prior *in vivo* observations (Muller et al., 2019). These results suggest that neither spatially segregated synaptic inputs nor dendritic compartmentalization are strictly required for differential control over broad and narrow spikes.

**Figure 4.**
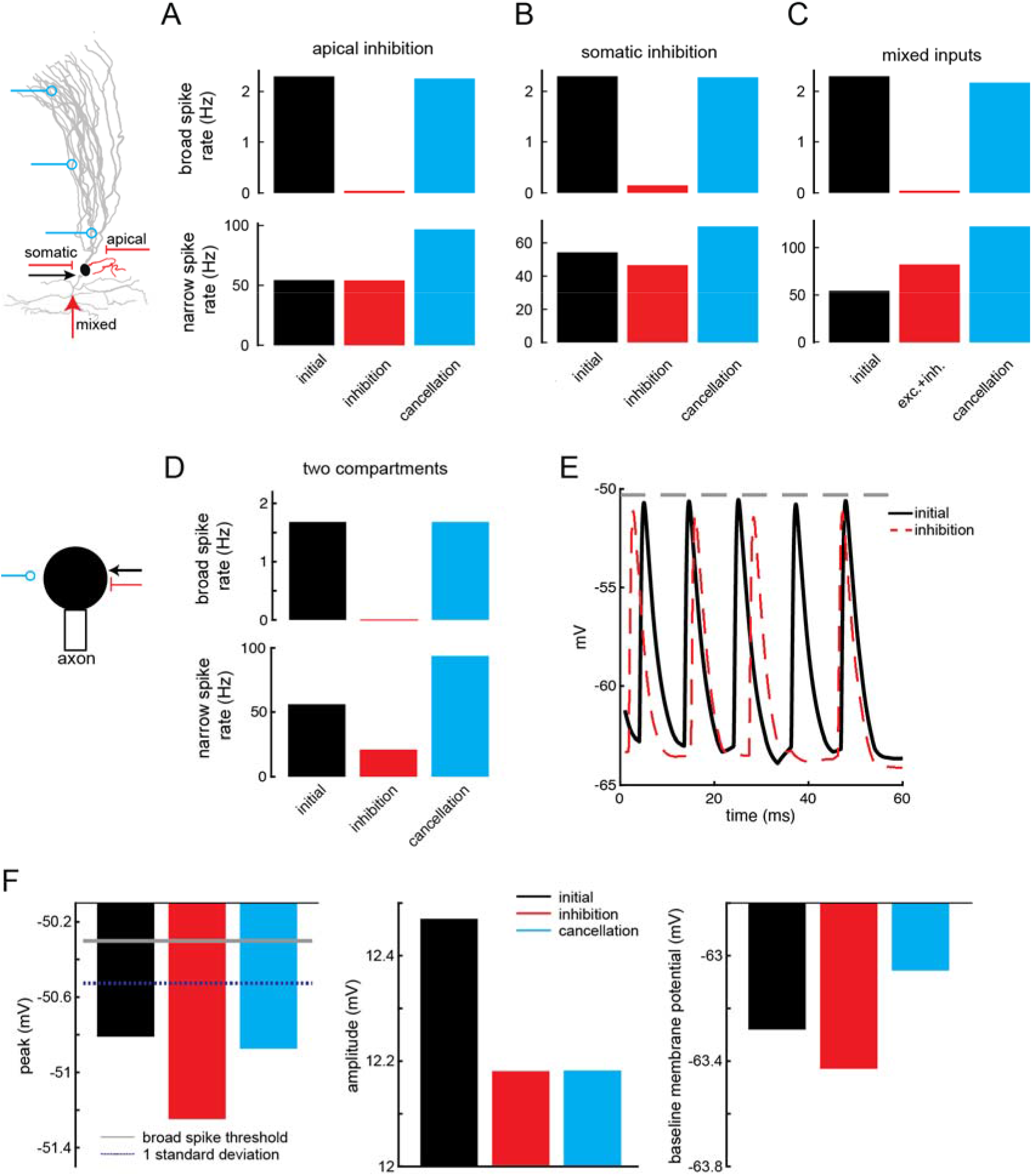
Negative image formation and transmission does not require dendritic compartmentalization. (**A-B**) Inhibition onto proximal apical dendrites (A) or soma (B) (indicated by the arrows, left) results in the formation and transmission of negative images in the model. (**C**) A mixture of excitatory and inhibitory inputs decreases the rate of broad spikes while increasing narrow spike firing. Cancellation of broad spike inhibition results in a further increase of narrow spike firing. (**D**) Formation and transmission of negative images can also be achieved in a simplified two-compartment model. (**E**) Trace from the two compartment model showing the reduction of backpropagating narrow spike amplitude by inhibitory input. (**F**) Mechanism of negative image formation and transmission in the two compartment model is the same as in the realistic model (cf. Figure 2B-D).

To test this further, we constructed a simple conductance-based integrate-and-fire model with only two compartments representing an axon and soma. Remarkably, the same qualitative results described for the morphologically realistic multi-compartment model were reproduced by the attenuation of the backpropagating axonal narrow spike in the somatic compartment (Figure 4D-F). While this result in no way excludes important functional roles for the numerous morphological, synaptic, and biophysical specializations of real MG cells, it suggests that the essential biophysical requirements for continual learning and signal transmission are surprisingly minimal.

## Discussion

Learning is typically associated with Hebbian forms of plasticity with homeostatic plasticity playing a stabilizing role by enforcing a return to equilibrium. Here we identified biophysical mechanisms that allow homeostatic synaptic plasticity to transmit learned signals. Using *in vivo* recordings and biophysical modeling we showed that sensory input modulates the rate of dendritic spikes by adjusting the amplitude of backpropagating axonal action potentials. Homeostatic plasticity counteracts these effects through changes in the underlying membrane potential, allowing the dendritic spike rate to be restored to equilibrium while simultaneously transmitting predictions through modulation of the axonal spike rate. The core requirements of the mechanism we describe--separate axonal and somatodendritic action potentials and an electronically distant site of axonal spike initiation—are found in many classes of neurons (Grace and Bunney, 1983; Hausser et al., 1995; Llinas et al., 1968; Spencer and Kandel, 1961; Spruston et al., 1995; Stuart and Sakmann, 1994), suggesting that roles for homeostatic plasticity in learning may be more widespread than is currently appreciated.

An action potential that arrives at the soma highly attenuated might seem an unlikely candidate for impacting dendrites. We find, to the contrary, that the small size of backpropagating axonal spikes in MG cells makes them susceptible to modulation and therefore an ideal candidate for flexibly controlling dendritic events. Importantly, the amplitude of backpropagating action potentials is highly sensitive to synaptic input (Llinas et al., 1968; Renshaw, 1942; Tsubokawa and Ross, 1996), much more sensitive than rates of action potential generation. This provides a mechanism for precise and, importantly, differential control of axonal and dendritic spikes that supports their separate functions. Whereas discussion of neuronal compartmentalization typically focuses on dendritic structure (London and Hausser, 2005; Major et al., 2013; Stuart and Spruston, 2015), our work provides a case in which the separation of the axon from the soma and dendrites is the essential element. While in our models this compartmentalization is based on a high resistance between axonal and somatic compartments, additional specializations (and potential sites of regulation) are likely to exist in real cells. For example, studies of medium superior olive neurons in the mammalian auditory brainstem provide evidence that the precise subcellular localization and inactivation properties of voltage gated sodium channels contribute to electrical isolation of the axon initial segment from the soma and dendrites (Ko et al., 2016; Scott et al., 2010).

Our results do not, of course, rule out important functions for MG cell dendrites or for additional biophysical specializations not included in our simplified models. Indeed, studies of a zone of the ELL involved in active electrolocation suggest that corollary discharge-driven inhibition of broad spikes is targeted to the putative site of broad spike initiation in the proximal apical dendrites (Sawtell et al., 2007) (Figure 3-figure supplement 3). Additional important questions for future studies are the anatomical basis for the dis-inhibitory circuit presumed to underlie sensory-evoked excitation of broad spikes in BS+ cells and the anatomical organization and functional role of recurrent connections between MG cells.

Both similarities and differences relevant to the present findings are found amongst the various vertebrate cerebellum-like structures. Homeostatic (anti-Hebbian) forms of plasticity at parallel fiber synapses are present in all cerebellum-like structures that have been thus far examined (Bell, 2002; Bell et al., 2008). In contrast, the presence of such plasticity at synapses onto the spiny apical dendrites of GABAergic Purkinje or Purkinje-like cells has only been described for the cerebellum, the mormyrid ELL, and the mammalian dorsal cochlear nucleus (DCN). In structures lacking Purkinje-like cells, such as the dorsal octavolateral nucleus of sharks and rays and the ELL of South American weakly electric, sensory cancellation may be a simpler one-stage process mediated by homeostatic plasticity at parallel fiber synapses onto glutamatergic output neurons (Bol et al., 2011; Nelson and Paulin, 1995). Cartwheel cells in the mammalian DCN, on the other hand, exhibit a number of similarities with MG cells, including firing distinct axonal and dendritic spikes (Kim and Trussell, 2007; Zhang and Oertel, 1993). While prior work has provided evidence for the cancellation of self-generated sounds in output cells of the DCN (Singla et al., 2017), possible roles for anti-Hebbian plasticity at parallel fiber synapses onto cartwheel cells remain to be investigated (Tzounopoulos et al., 2004). The possible computational advantages of performing sensory cancellation in two-stages, as opposed to one, also remain to be elucidated.

Roles for homeostatic plasticity in transmitting learned signals may extend beyond cerebellum-like structures. Anti-Hebbian spike timing-dependent plasticity, similar to that at granule-MG cell synapses, has been documented at synapses in the striatum (Perez et al., 2022) and neocortex (Letzkus et al., 2006; Ruan et al., 2014). While homeostatic plasticity of inhibitory synapses onto pyramidal cells has been hypothesized to underlie responses to prediction errors in sensory cortical neurons (Hertag and Sprekeler, 2020; Keller and Mrsic-Flogel, 2018), less is known about the mechanisms for transmitting predictions between cortical layers or regions. MG cells play an analogous role by transmitting learned predictions across processing stages of the ELL. Many cortical neurons fire distinct axonal and dendritic spikes, suggesting the possibility that mechanism similar to those described here may allow homeostatic plasticity to contribute to predictive processing in the cerebral cortex.

## Materials and methods

### Experimental model and subject details

Male and female Mormyrid fish (7-12 cm in length) of the species *Gnathonemus petersii* were used in these experiments. Fish were housed in 60 gallon tanks in groups of 5-20. Water conductivity was maintained between 40-65 microsiemens. All experiments performed in this study adhere to the American Physiological Society’s *Guiding Principles in the Care and Use of Animals* and were approved by the Institutional Animal Care and Use Committee of Columbia University.

For surgery to expose the brain for recording, fish were anesthetized (MS:222, 1:25,000) and held against a foam pad. Skin on the dorsal surface of the head was removed and a long-lasting local anesthetic (0.75% Bupivacaine) was applied to the wound margins. A plastic rod was cemented to the anterior portion of the skull to secure the head. The posterior portion of the skull overlying the ELL was removed and the valvula cerebelli was reflected laterally to expose the eminentia granularis posterior (EGp) and the molecular layer of the ELL, facilitating whole-cell recordings from the ventrolateral zone of the ELL. Gallamine triethiodide (Flaxedil) was given at the end of the surgery (~20 μg/cm of body length) and the anesthetic was removed. Aerated water was passed over the fish’s gills for respiration. Paralysis blocks the effect of electromotoneurons on the electric organ, preventing the EOD, but the motor command signal that would normally elicit an EOD continues to be emitted at a rate of 2 to 5 Hz.

### Electrophysiology

The EOD motor command signal was recorded with a Ag-AgCl electrode placed over the electric organ. The command signal is the synchronized volley of electromotoneurons that would normally elicit an EOD in the absence of neuromuscular blockade. The command signal lasts about 3 ms and consists of a small negative wave followed by three larger biphasic waves. Onset of EOD command was defined as the negative peak of the first large biphasic wave in the command signal. For pairing experiments, the EOD mimic was presented 4.5 ms following EOD command onset. Recordings were started ~1 hour after paralysis.

Methods for *in vivo* whole-cell recordings were the same as in prior studies of the mormyrid ELL (Muller et al., 2019; Sawtell, 2010). Briefly, electrodes (8-15 MΩ) were filled with an internal solution containing, in mM: K-gluconate (122); KCl (7); HEPES (10); Na2GTP (0.4); MgATP (4); EGTA (0.5), and 0.5-1% biocytin (pH 7.2, 280-290 mOsm). No correction was made for liquid junction potentials. Membrane potentials were recorded and filtered at 10 kHz (Axoclamp 2B amplifier, Axon Instruments) and digitized at 20 kHz (CED micro1401 hardware and Spike2 software; Cambridge Electronics Design, Cambridge, UK). Only cells with stable membrane potentials more hyperpolarized than −40 mV and broad spike amplitudes >40 mV were analyzed. In contrast to broad spikes, narrow spike amplitude varied across recordings from ~15 mV (similar to values obtained from somatic recordings *in vitro*) to indistinguishable from subthreshold synaptic events. The latter, which were typically obtained at more superficial recording depths corresponding to the ELL molecular layer, were classified as putative apical dendritic recordings (see figure supplement 1D).

### Electrosensory stimulation

The EOD mimic was a 0.2 ms duration square pulse delivered between an electrode in the stomach and another positioned near the electric organ in the tail. The amplitude was 25-50 μA at the output of the stimulus isolation unit (stomach electrode negative). Recordings from ampullary afferents showed that firing rate modulations evoked by this mimic are within the range of those induced by the fish’s natural EOD (Bell and Russell, 1978). We use the terms sensory input or sensory response to refer to the effect of the mimicked electric field on the ELL. Because we do not include prey-like electric fields the sensory input we discuss is entirely predictable on the basis of the EOD command signal and is therefore entirely uninformative to and ‘unwanted’ by the fish. Thus, we consider a situation where the ELL attempts to cancel *all* of its sensory input. It is important to appreciate that, in a natural setting, the mechanisms we analyze would only cancel the predictable self-generated component of the sensory input, leaving the unpredictable inputs of interest to the fish intact. To isolate responses to sensory versus corollary discharge we analyzed periods in which sensory stimuli were delivered independent of the EOD motor command. In some cases, sensory responses were isolated from periods in which the sensory stimuli were paired with the EOD motor command by off-line subtraction of responses to the EOD motor command alone.

### Quantification and Statistical Analysis

Data were analyzed off-line using Spike2 (Cambridge Electronic Design) and custom Matlab code (Mathworks, Natick, MA). Biophysical model analysis was performed using custom Python3 code. Non-parametric tests were used for testing statistical significance. Unless otherwise indicated, we used the two-sided Wilcoxon rank sum test for unpaired samples and the Wilcoxon signed ranks test for paired samples. Differences were considered significant at *P* < 0.05. 3 stars indicate a *P* < 0.001.

### Biophysical model

The compartmental model was based on a morphological reconstructed MG cell and consisted of 78 compartments further divided to 230 segments (Muller et al., 2019). Simulation of cell activity was done using NEURON software and Python 3 wrapper (Carnevale and Hines, 2006). Voltage gated Na+ and K+ channels inserted in the apical dendrites and axon are Hodgkin-Huxley type channels. Temperature was set to 20° Celsius. The attenuation of axonal spikes in the model arises simply due to the resistance between axonal and somatodendritic compartments. Voltage-gated channel conductances were adjusted (see **Table 1**) to achieve the higher spike threshold for broad versus narrow spikes that is observed experimentally.

**Table 1.** Values of biophysical parameters for the different compartments. ‘Rest’ includes the soma, the somatic-connected apical compartment and all basal dendrite compartments. *g_l_*, is leakage conductance. 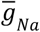, and 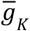 are the maximal conductances of the sodium and potassium channels, respectively.

| | $g_l$<br>(S/cm<sup>2</sup>) | leakage<br>reversal<br>potential<br>(mV) | axial<br>resistance<br>(Ωcm) | capacitance<br>(μF/cm<sup>2</sup>) | $\bar{g}_{Na}$<br>(S/cm<sup>2</sup>) | $\bar{g}_K$<br>(S/cm<sup>2</sup>) |
| --- | --- | --- | --- | --- | --- | --- |
| axon | 0.0003 | -65 | 100 | 1 | 4 | 0.5 |
| AIS | 0.0003 | -65 | 100 | 1 | 0.168 | 0.05 |
| apical | 0.0003 | -65 | 100 | 1 | 0.1 | 0.008 |
| rest | 0.0003 | -65 | 100 | 1 | 0 | 0 |

To drive baseline spiking in the model cell (the condition we term *initial*), we injected Gaussian current noise into the soma (0.5 ms timesteps) with a standard deviation chosen to evoke ~50 Hz narrow spike firing and ~2 Hz broad spike firing. To drive excitatory and inhibitory responses we added AMPA and GabaA synaptic channels (Destexhe et al., 1994). Reversal potential of the AMPA and GabaA channel are 0 mV and −65 mV, respectively. The AMPA excitatory input was inserted into all apical dendrite compartments (49 compartments 175 segments). The AMPA and GabaA inputs were constant, in which each relevant compartment received a synaptic input with timing onset 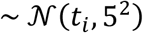 where *t_i_* = 10 · *i*. For basal dendrites inhibition (21 compartments (45 segments), conductance was 0.1 μS and excitatory conductance was 7.65e-5 μS. For somatic inhibition, conductance was 0.04 μS and excitatory conductance was 1.85e-5 μS. For apical inhibition, conductance was 0.01 μS and excitatory conductance was 7.1e-5 μS. Apical inhibition was inserted into proximal apical compartments (11 compartments (19 segments)) defined as those whose center is within 100 *μm* of the center of the soma.

### Two compartment model

Conductance based integrate-and-fire model was used for the two compartment model (Fig 4d). The equations for somatic and axonal membrane potential are:

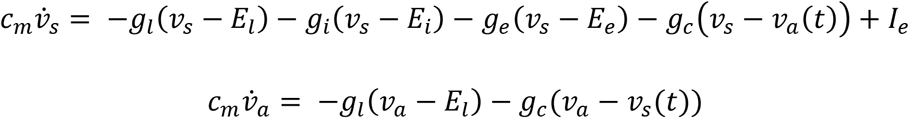

Where *g_l_* is the leakage conductance, *g_c_* is the intercompartment conductance and *g_i_* and *g_e_* are the inhibitory and excitatory conductances respectively. *I_e_* is external current (with Gaussian noise) and is set to produce ~50 Hz narrow spike and ~2 Hz broad spike rates. When the axon reaches the threshold for axonal spike, a spike shape plus a refractory period is imposed in the axon. Broad spike rate was determined by the number of times the backpropagating axonal spike reached a high threshold in the soma (this threshold was defined as the 97th percentile of the backpropagating spike-peak in the *initial* period).

### Measuring narrow spike amplitude differences

Quantifying narrow spike amplitude differences induced by sensory input is complicated by the strong dependence of narrow spike amplitude on baseline membrane potential observed *in vivo* (negative slope in Figure 1I). To account for this effect, we fit the slope of the relationship between narrow spike amplitude and baseline membrane potential and report the difference across conditions in the bias of these slopes. Similarly, to measure difference between expected and actual amplitude (figure supplement 6A) we first fit a slope to the relationship between amplitude and baseline membrane potential and then measure the distance from the fit.

We hypothesize (figure supplement 1H-I) that the attenuation of backpropagating narrow spike amplitude is linearly proportional to the amplitude:

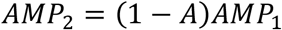

Then if we divide by average recorded mean we have the following equality:

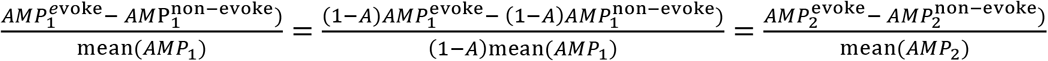

The average narrow spike amplitude differs widely across recordings (see figure supplement 1D), presumably due to recording location in the soma versus the proximal apical dendrites. Hence, to compare differences in narrow spike amplitude evoked by sensory stimuli across recordings we report the percentage change in narrow spike amplitude relative to the average narrow spike amplitude for each cell. The same reasoning applies to analysis of the relationship between amplitude and baseline membrane potential across different cells (figure supplement 7A).

### F-I curve for narrow spikes

The fit between membrane potential and spike rate is approximately linear (see example in Figure 3E). To minimize the effect of outliers we quantify the change in rate/mV as:

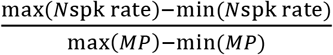

### Measuring effect of sensory input on amplitude

Recorded amplitude of *in vivo* narrow spikes may change over the course of the recording (as the quality of the recording changes). Therefore, we meausred amplitude differences in relation to a control window within same recording period. Changes in amplitude from the first to the second half of the recorded period (Figure 3d) was measured as the change in amplitude relative to the control window within each half of the recording.

## Data and code availability

Model code is available at: http://modeldb.yale.edu/267596, password: abbottsawtell Data and data code is available at: https://datadryad.org/stash/share/5oEoH42oOWfSQ07fLc3mMIa92CdzrXE8wYSKvvV-HhE

## Acknowledgements

This work was supported by grants from the NIH (NS075023) and Irma T. Hirschl Trust to N.B.S., by a grant from the NIH (NS118448) to N.B.S. and L.F.A., and by a grant from the Swartz Foundation to S.Z.M. L.F.A. was further supported by the Gatsby and Simons Foundations and by NSF NeuroNex Award DBI-1707398.

Zuckerman Mind Brain Behavior Institute, Department of Neuroscience, Columbia University, New York, NY 10027

Salomon Z. Muller, L.F. Abbott and Nathaniel B. Sawtell

Department of Physiology and Cellular Biophysics, Columbia University, New York, NY 10027 L.F. Abbott

## Contributions

N.B.S., S.Z.M., and L.F.A. conceived of the project and designed the experiments. N.B.S. performed the experiments. S.Z.M. and N.B.S. analyzed the data. S.Z.M. and L.F.A. performed the modeling. N.B.S., L.F.A. and S.Z.M. wrote the manuscript.

## Competing financial interests

The authors declare no competing financial interests

---

**Figure 1-figure supplement 1.**
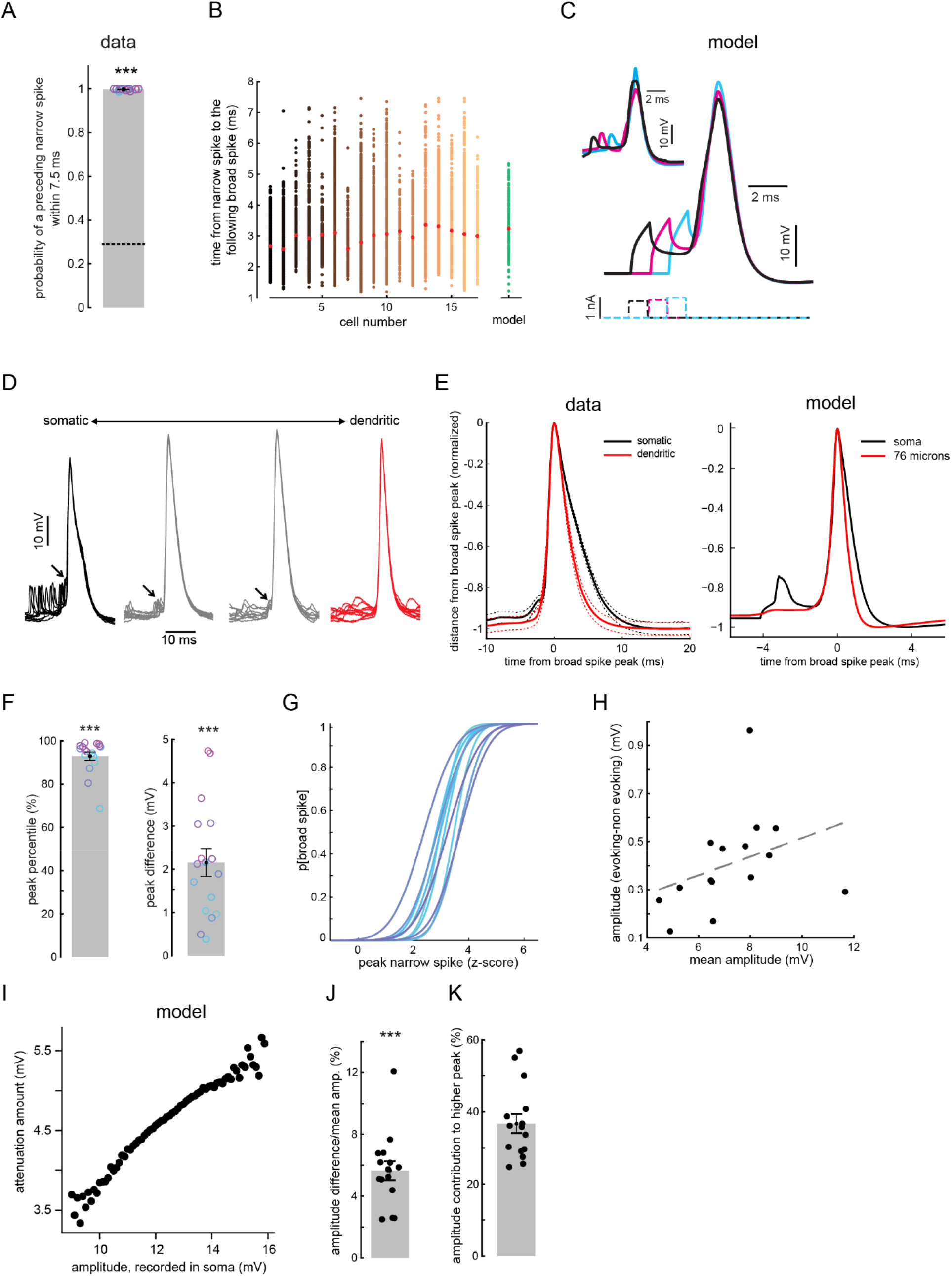
Narrow spikes contribute to evoking broad spikes. (**A**) Broad spikes are nearly always preceded by a narrow spike within a brief interval. Average chance level is indicated by the dashed line (n=17, p < 0.001). (**B**) Distribution of times from the peak of a preceding narrow spike to the peak of the following broad spike in data (n = 17) versus the model. Red star denotes the mean. (**C**) Injecting a brief depolarizing current into the model soma (with active conductances in the axon turned off) evokes broad spikes at similar delays to those observed between narrow and broad spikes (inset). The latency of the evoked broad spike is inversely proportional to the strength of the depolarizing current (bottom dashed lines). (**D**) Overlaid traces aligned to occurrence of a broad spike for putative somatic (left) versus dendritic (right) MG cell recordings. Recordings with narrow spikes ≥4 mV were classified as putative somatic (left) and those with narrow spikes not clearly indistinguishable from synaptic potentials were classified as putative dendritic (right). Many recordings exhibited narrow spikes with intermediate amplitudes, as expected for a passively backpropagating axonal spike. Narrow spikes preceded broad spikes (arrow) in all cases in which they were detectable. In dendritic recordings broad spikes often arose directly from the underlying membrane potential, similar to late firing dendritic branches in the model (*green trace* in Figure 1F). (**E**) Broad spike waveforms are narrower in putative dendritic versus putative somatic recordings (width at half height: 3.1 ms, n = 11 for dendritic versus 4.1 ms, n = 17 for somatic recordings, p<0.001). Dashed lines indicate SEM. Broad spikes were also wider in the soma in the model (right) due to summation of broad spikes from multiple apical dendritic branches. (**F**) The membrane potential peak reached by narrow spikes preceding broad spikes is in the top 10 percentile of all narrow spikes (left bar, n=17, p<0.001), and the average peak difference between evoking and non-evoking narrow spikes is ~2.2mV (right bar, n=17, p<0.001). (**G**) Preceding narrow spike peak strongly predicts probability of evoking a broad spike forming a typical logistic curve (n=10). (**H**) The size of the difference between the amplitude of narrow spikes preceding broad spikes and non-preceding narrow spikes depends on average recorded narrow spike amplitude (n=15). (**I**) In the model, spatial attenuation of the backpropgating narrow spike depends on the amplitude recorded in the soma, with larger spikes exhibiting greater attenuation. This explains why average recorded narrow spike amplitude affects the size of the amplitude difference between preceding and non-preceding narrow spikes (see Materials and methods). (**J**) Amplitude of narrow spikes preceding broad spikes are larger than non-preceding narrow spikes by ~5.5% (n=15, p<0.001) (see Materials and methods as to why we use this measure). (K) The more depolarized peak of preceding narrow spikes is due both to larger amplitude and to a more depolarized underlying membrane potential (Figure 1I), with amplitude contributing to ~35% of the difference (n=15).

**Figure 2-figure supplement 1.**
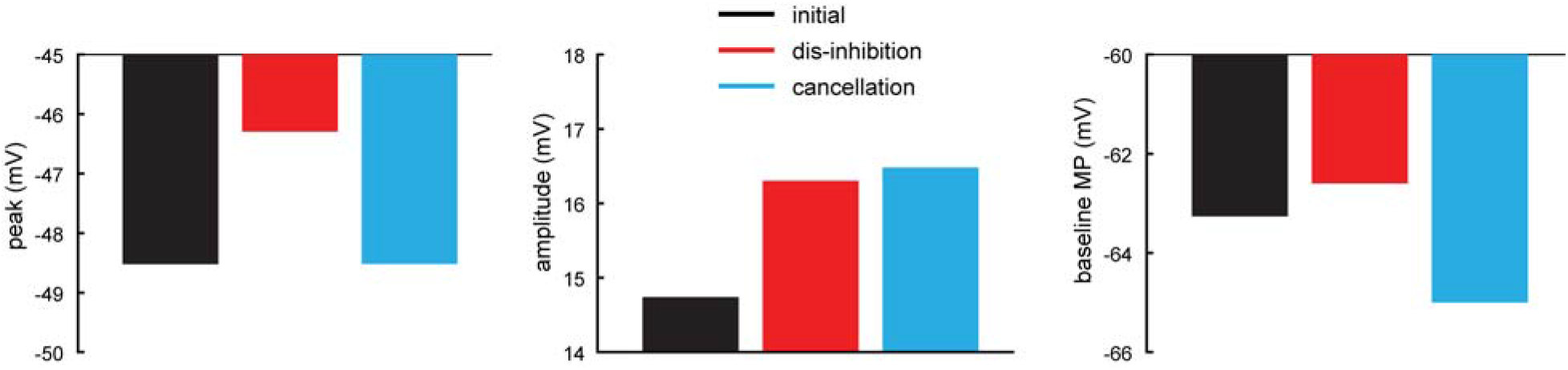
Formation and transmission of negative images in the BS+ MG cell sub-type. Analytical results (see Appendix 1) for BS+ in which dis-inhibition increases broad spike rate. Here, disinhibition leads to higher peak of narrow spike, mostly due to increase in narrow spike amplitude. Removal of excitatory input (cancellation) restores narrow spike peak to initial values. The restoration is not done by reversing the effect of disinhibition on narrow spike amplitude, but rather by a hyperpolarization of the baseline membrane potential.

**Figure 2-figure supplement 2.**
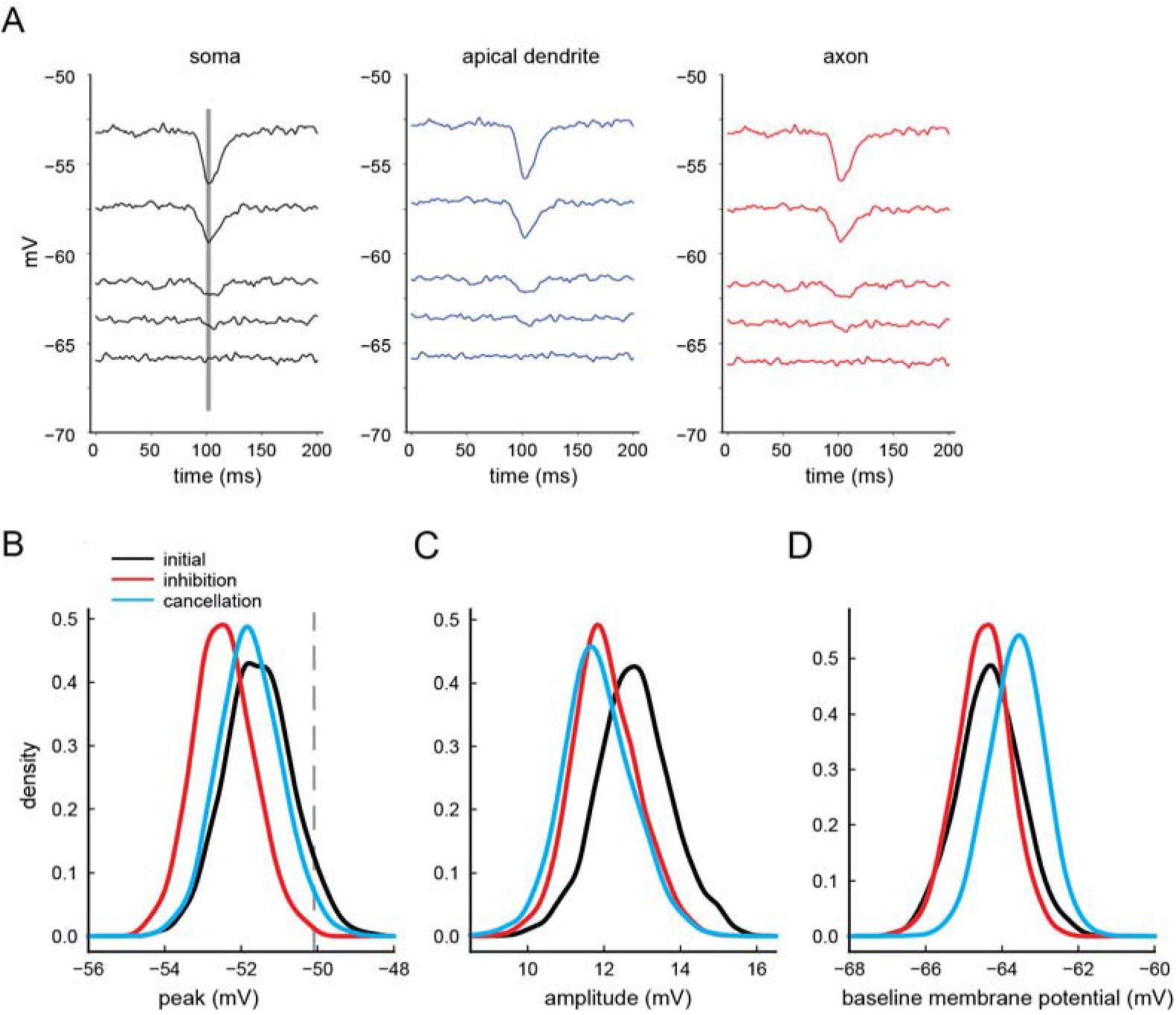
Biophysical explanation for negative image formation and transmission in the model. (**A**) As expected, hyperpolarization due to a transient increase in inhibitory current (line) increases when the baseline membrane potential is further from the reversal potential for inhibitory input (−65mV). Importantly, the effect of inhibition is similar across different model compartments, confirming that inhibitory input has a minimal effect on narrow spike rate because narrow spike threshold is close to the reversal potential for inhibitory input. In this simulation all active conductances were turned off to clearly see effect of inhibition on the membrane potential. (**B-D**) Similar to Figure 2B-D, but here we plot the entire distribution of narrow spike peak (B), narrow spike amplitude (**c**) and baseline membrane potential (D) values for the different input conditions. Gray line in B indicates broad spike threshold.

**Figure 2-figure supplement 3.**
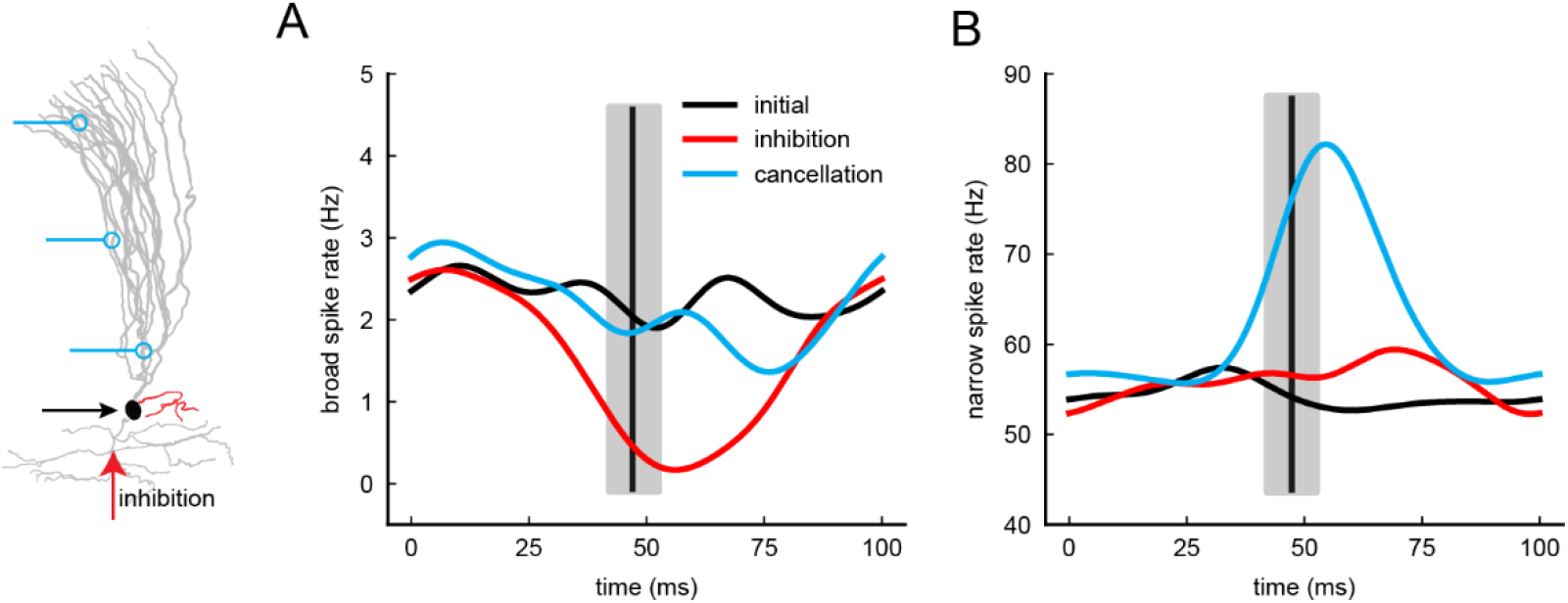
Formation and transmission of a time-varying negative image in the model. (**A-B**) Similar to Figure 2A but here the inhibitory and excitatory inputs are transient (vertical lines denote average timing onset of synaptic input and gray bars denote the standard deviation). This mimics *in vivo* conditions where predictable electrosensory input is evoked by the fish’s electric organ discharge pulse. Inhibitory input onto basal dendrites can selectively inhibit broad spikes (red). Adding excitatory inputs onto apical dendrites to simulate the process of negative image formation reduces the effects of inhibition on broad spike rate (cyan) and results in an increase in narrow spike firing with a temporal profile opposite to the effect of the sensory input on broad spikes, i.e. the transmission of a negative image of the effects of the sensory input on broad spike firing.

**Figure 2-figure supplement 4.**
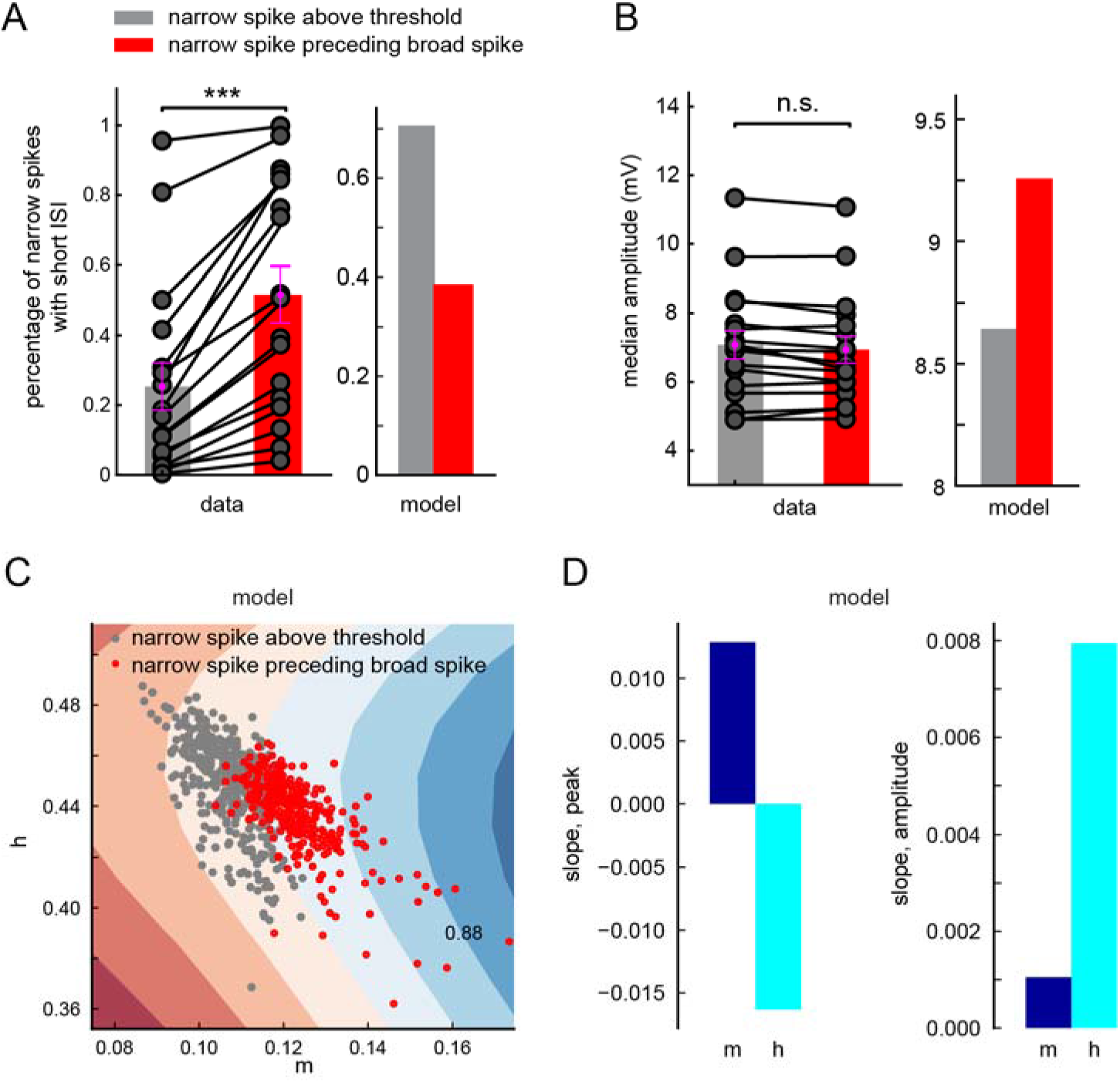
The probability of a narrow spike evoking a broad spike also depends on prior history of activity. (**A**) In recorded MG cells, the probability of a narrow spike evoking a broad spike is increased when it is preceded within a short interval by another narrow spike at a brief inter-spike interval (ISI) (n = 17, p < 0.001). ISI values were chosen individually for each cell depending on the narrow spike rate. The gray bar represents narrow spikes that exceeded broad spike threshold (defined as the 10th percentile of narrow spike peaks preceding a broad spike) but nevertheless failed to evoke a broad spike. The opposite effect was observed in the model; short narrow spike ISIs decreased the probability of evoking a broad spike in the model. This decrease is an expected consequence of the Hodgkin-Huxley type voltage-gated channels used in the model. (**B**) For narrow spikes that exceeded broad spike threshold, there was no difference in amplitude between preceding and non-preceding narrow spikes (n = 17, p = 0.37). A different effect was observed in the model; narrow spike preceding broad spikes have on average larger amplitude. This is an expected consequence of the Hodgkin-Huxley type voltage-gated channels used in the model (as we show in C-D). (**C**) Values for sodium channel activation (m), sodium channel inactivation (h) and potassium channel activation (n) were measured 0.65ms after narrow spike peak in a compartment near the site of broad spike initiation in the model. Using a support vector machine we could largely separate (with accuracy of 88%) the broad spike-evoking (red dots) and non-evoking (gray dots) narrow spikes using just m and h values (we ignore n here since it is highly correlated with h). Colored bands denote degrees of separation-confidence, with darker colors denoting higher confidence. (**D**) While the peak of the narrow spike is correlated with both m and h, it is h that is sensitive to amplitude because it strongly depends on the recent history of the membrane potential, with a rapid rise to threshold increasing spike probability.

**Figure 3-figure supplement 1.**
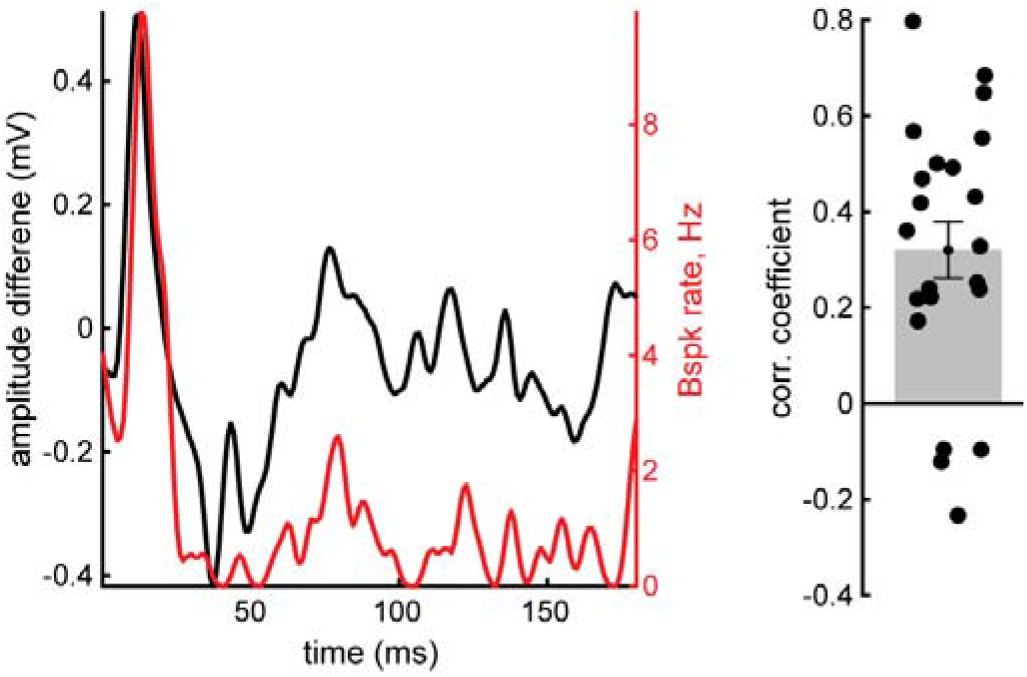
Correlation between sensory input and narrow spike amplitude *in vivo*. **Left**, example traces illustrating broad spike response to an electrosensory stimulus (red) and the difference between the measured narrow spike amplitude and that expected given the baseline membrane potential (amplitude difference, see Materials and methods). **Right**, summary across cells of the correlation between broad spike rate and amplitude difference (r = 0.35, n = 22 stimulus periods from 16 cells). Multiple stimulus polarities and amplitudes were tested in some cells.

**Figure 3-figure supplement 2.**
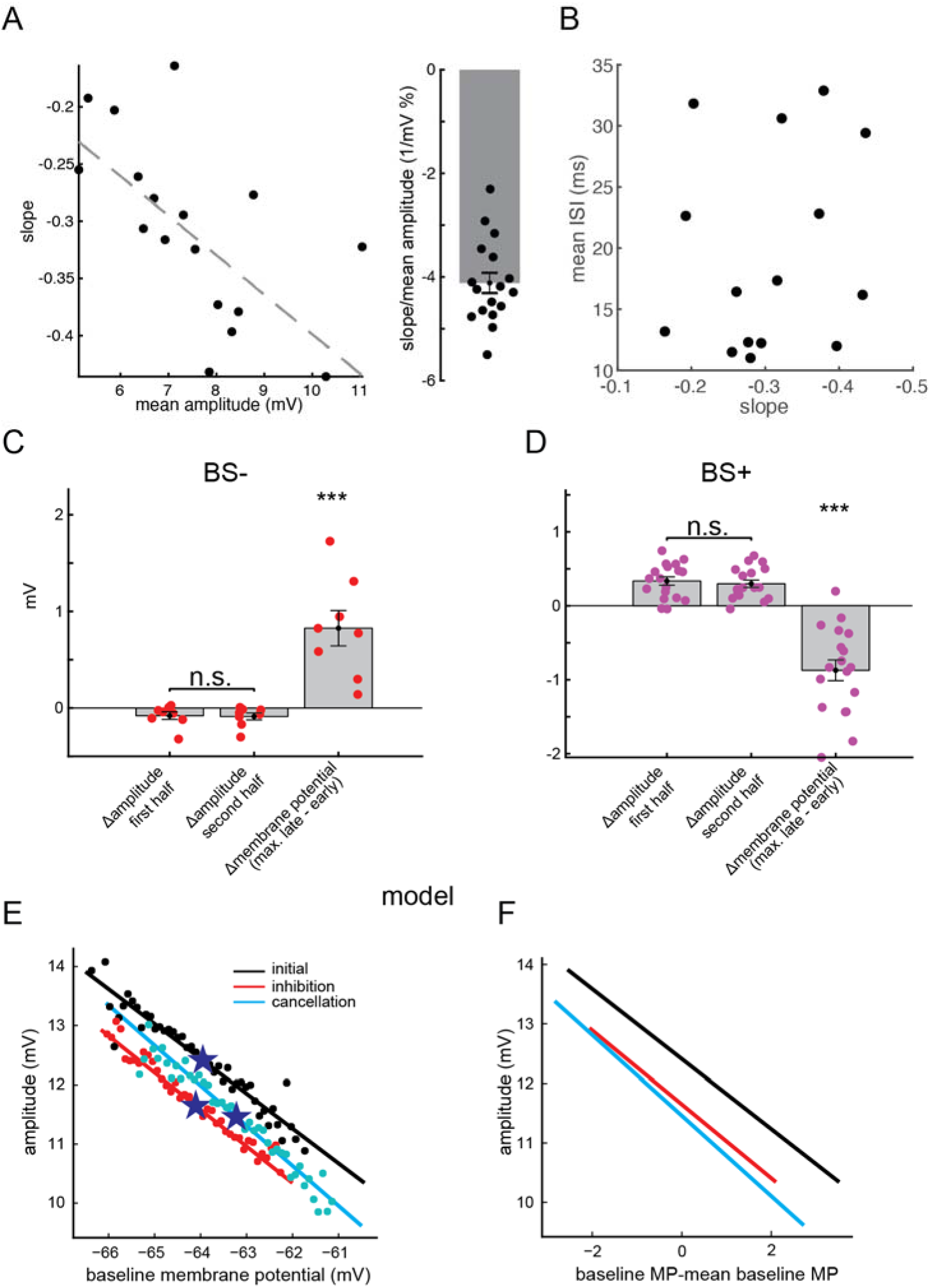
Effects of the baseline membrane potential on narrow spike amplitude. (**A**) Left, the slope of the relationship between narrow spike amplitude and baseline membrane potential depends on the average narrow spike amplitude. The latter varies across recordings, likely due to recording location in the soma versus proximal apical dendrites. The reduction in slope is consistent with the observation in the model that spatial attenuation magnitude is proportional to narrow spike amplitude (figure supplement 1I). Right, the average slope as a percentage of spike size is 4.1% (n = 16) (see Materials and methods as to why we use this measure). (**B**) The slope of the relationship between narrow spike amplitude and baseline membrane potential is independent of the mean inter-spike interval, suggesting that the change in amplitude is independent of spiking history. (**C-D**), In the main text we show that learning to cancel the effect of sensory input (the period we call here ‘second half’) changes the underlying membrane potential. This results in average amplitude change due to the slope of the relationship between narrow spike amplitude and baseline membrane potential (Figure 3D). However, for spikes arising from the same baseline membrane potential, the difference in amplitude between sensory evoked and control windows (Figure 3A-C) is not affected by learning. (**E-F**), In the model, cancellation (i.e. the ‘second half’) does change the amplitude for spikes arising from same baseline membrane potential (E, the stars in E denote mean baseline membrane potential) but the average amplitude is minimally changed by learning (F). As a result, the amplification of negative images observed *in vivo* is absent in the model. Whereas the slope of the relationship between narrow spike amplitude and baseline membrane potential in the model is due to deviations from equilibrium driven by fluctuations in the baseline current inputs.

**Figure 3-figure supplement 3.**
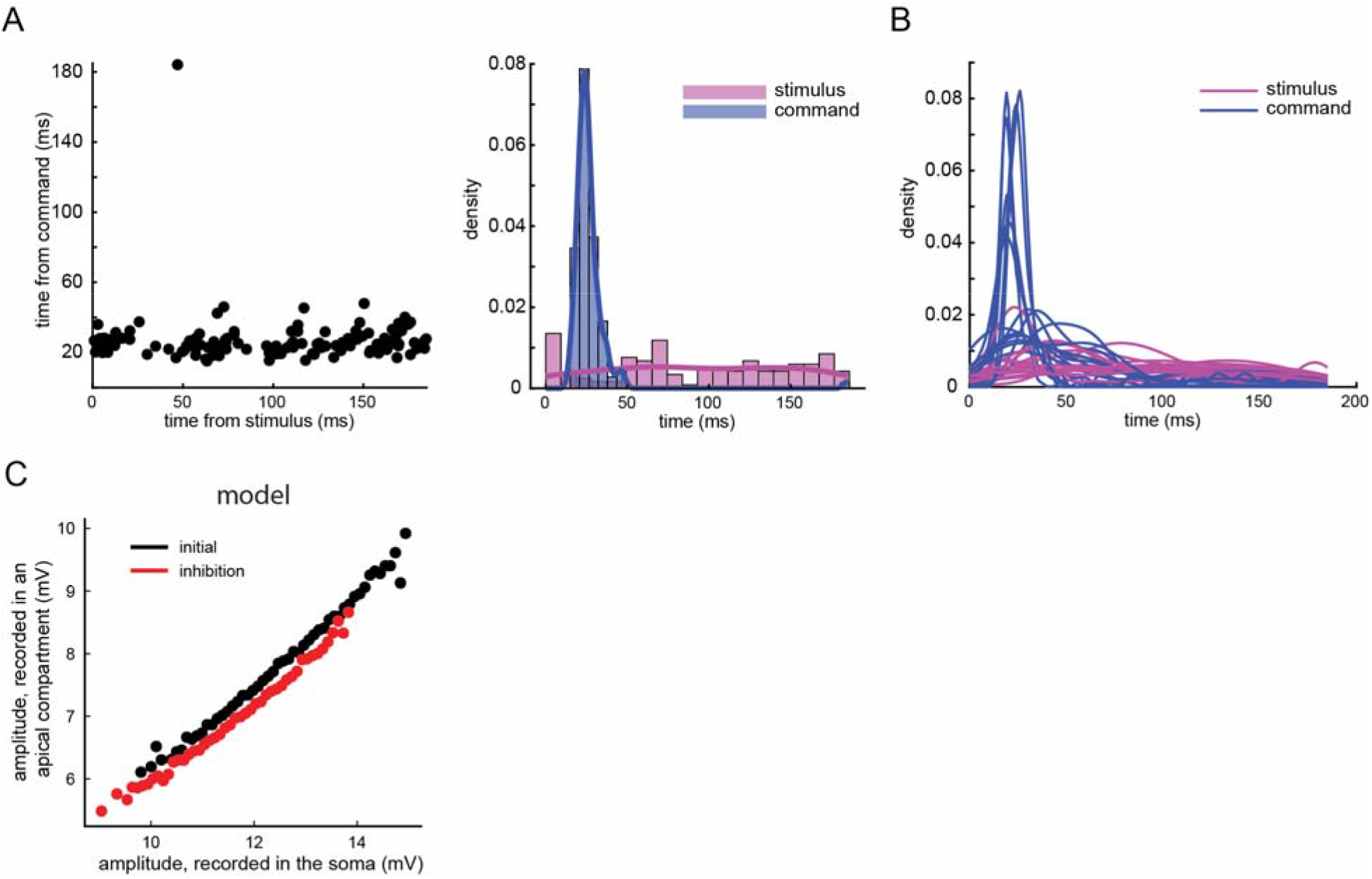
Additional effects of corollary discharge input on broad spike firing may not be visible at the soma. Non-evoking narrow spikes that cross threshold for evoking a broad spike in somatic recordings (threshold is calculated here as the 20th percentile peak of the broad spike-evoking narrow spikes), tend to systematically occur ~20-40 ms after the EOD motor command, corresponding to the period of peak corollary discharge-evoked responses in MG cells. This analysis was performed for periods during which an electrosensory stimulus was delivered independent of the EOD motor command. (**A**) Example from one period shown as a scatter plot (left, each dot is a non-evoking narrow spike that crossed threshold) and a histogram (right). (**B**) Same analysis as in B. Each line is a different cell (n=12). (**C**) Model analysis showing that the effect of inhibition on reducing narrow spike amplitude continues to grow as the spike continues to backpropagate even when inhibition is inserted in the basal dendrites. This observation is consistent with the possibility that effects of inhibition onto apical dendrites may not be visible in the soma.

## Notes

### Competing Interest Statement

The authors have declared no competing interest.

